# lncRNA *3222401L13Rik*/*ENSG00000272070* modulates microglial inflammatory programs in association with PU.1

**DOI:** 10.1101/2025.10.31.685723

**Authors:** Ranjit Pradhan, M Sadman Sakib, Lalit Kaurani, Dennis M. Krüger, Tonatiuh Pena, Susanne Burkhardt, Anna-Lena Schütz, Deborah Kronenberg-Versteeg, Ivana Delalle, Farahnaz Sananbenesi, Andre Fischer

**Affiliations:** Department for Systems Medicine and Epigenetics, German Center for Neurodegenerative Diseases (DZNE), Göttingen, Germany; Bioinformatics Unit, German Center for Neurodegenerative Diseases (DZNE), Göttingen, Germany; Research Group for Genome Dynamics in Brain Diseases, German Center for Neurodegenerative Diseases, Göttingen, Germany; Department of Cellular Neurology, Hertie Institute for Clinical Brain Research, University of Tübingen, Germany; Germany and German Center for Neurodegenerative Diseases (DZNE) Tübingen, Germany; Department of Pathology and Laboratory Medicine, Boston University School of Medicine, 670 Albany Street, Boston, MA 02118, USA; Cluster of Excellence “Multiscale Bioimaging: from Molecular Machines to Networks of Excitable Cells” (MBExC), University of Göttingen, Germany; Department of Psychiatry and Psychotherapy, University Medical Center Göttingen, Göttingen, Germany

**Keywords:** long non-coding RNA (lncRNA), non-coding RNAome, microglia, PU.1 (SPI1), neuroinflammation, Alzheimer’s disease

## Abstract

Long non-coding RNAs (lncRNAs) are emerging as key regulators of brain function, but their contribution to microglial aging and neurodegenerative disease remains largely unknown. Because only 1.5% of the human genome encodes proteins, whereas the vast majority of transcripts belong to the largely unexplored non-coding RNAome, elucidating the functions of non-coding RNAs provides an unprecedented opportunity to expand the space for therapeutic discovery. We recently identified the glia-enriched lncRNA *3222401L13Rik* as upregulated in the aging mouse hippocampus. Here, we investigated its function in microglia and its human homolog *ENSG00000272070*. We found that *3222401L13Rik* is expressed in both astrocytes and microglia and increases with age. Knockdown of *3222401L13Rik* in primary microglia led to enhanced expression of pro-inflammatory cytokines, including TNFα, and increased phagocytic activity. RNA-sequencing revealed widespread transcriptional changes enriched for TNF and complement signaling pathways. The human homolog *ENSG00000272070* showed conserved functions in iPSC-derived microglia, where its loss similarly promoted inflammatory gene expression and phagocytosis. Mechanistically, *3222401L13Rik* interacts with the microglial transcription factor PU.1, and its depletion overlapped with PU.1-driven transcriptional programs. Consistent with these findings, *ENSG00000272070* expression was significantly reduced in postmortem Alzheimer’s disease (AD) brains, and AD-associated genes were enriched among *3222401L13Rik*-regulated targets. Together, our results identify *3222401L13Rik*/*ENSG00000272070* as a conserved, aging-associated lncRNA that modulates microglial inflammatory states through interaction with PU.1. This work links glial lncRNA regulation to AD-related neuroinflammation and suggests *3222401L13Rik* as a potential molecular target to fine-tune microglial activity in neurodegenerative diseases.

**Highlights:**

- *3222401L13Rik* is a glia-enriched long non-coding RNA regulating microglial state
- Knockdown of *3222401L13Rik* increases TNFα signaling
- The human homolog *ENSG00000272070* shows reduced expression in AD brains
- *3222401L13Rik* interacts with the AD-associated transcription factor PU.1
- A conserved *3222401L13Rik*–PU.1 axis modulates microglial inflammation in aging and disease

## Introduction

Microglia are the resident immune cells of the central nervous system, where they perform immune surveillance, remove cellular debris, and support neuronal networks through maintenance of synapses and homeostatic signaling {Prinz, 2014} {Colonna, 2017}. During aging, microglia undergo profound molecular and functional alterations that shift them toward a chronically activated, dysfunctional state {Damani, 2011}. These changes are increasingly recognized as drivers of neurodegenerative diseases such as Alzheimer’s disease (AD), Parkinson’s disease (PD), and frontotemporal dementia (FTD) {Davies, 2016} {Streit, 2009}. Aged microglia show reduced motility and surveillance capacity {Costa,, 2021} and decreased ability to clear pathogenic proteins such as amyloid-β (Aβ) and α-synuclein {Floden, 2011} {Bido, 2021} {Scheffold, 2016}. Although transcriptomic studies have revealed extensive age-related reprogramming of microglia {Grabert, 2016} {Soreq, 2017}, the underlying regulatory mechanisms remain poorly understood.

Long non-coding RNAs (lncRNAs) have emerged as key regulators of gene expression and cellular identity in the brain {Statello, 2021} {Mattick, 2023}. Defined as transcripts exceeding 300 nucleotides without protein-coding potential, lncRNAs can act as molecular scaffolds, guides, or decoys to modulate transcriptional and post-transcriptional processes {Quinn,, 2015}. Approximately 40 % of annotated human lncRNAs show brain-specific expression {Derrien, 2012}, highlighting their importance in neural cell-type specialization and disease. Despite their abundance, the role of lncRNAs in microglial aging and neurodegeneration has remained largely unexplored.

In our previous study {Schröder,, 2025}, we identified *3222401L13Rik* as a glia-enriched lncRNA that is upregulated in the aging mouse hippocampus, with a conserved human homolog *ENSG00000272070*. We demonstrated that in astrocytes, *3222401L13Rik* regulates neuronal support and synaptic organization through interaction with the transcription factor Neuronal PAS Domain Protein 3 (Npas3), thereby contributing to adaptive mechanisms that may mitigate age-related synaptic decline. At that time, *3222401L13Rik* had not been studied in any biological context {Schröder, 2025}.

Interestingly, *3222401L13Rik* is not restricted to astrocytes but is also expressed in microglia. However, its role in microglial biology has never been investigated. Given that microglia are central to neuroinflammatory and neurodegenerative processes, we sought to determine how *3222401L13Rik* influences microglial function and whether its regulatory network overlaps with pathways implicated in AD.

Here, we show that *3222401L13Rik* and its human homologue *ENSG00000272070* fine-tunes microglial transcriptional programs and inflammatory activity. Using primary microglia and human iPSC-derived microglia, we demonstrate that loss of this lncRNA increases *TNFα* expression, enhances phagocytic activity, and alters neuronal network function in microglia– neuron co-cultures. We further show that *3222401L13Rik* interacts with the transcription factor PU.1, a master regulator of microglial identity {Yeh, 2019} {Saeki, 2025}, and that its human homolog is downregulated in AD brains and co-regulates a subset of AD-associated genes.

Together, these findings extend our previous work and establish *3222401L13Rik*/*ENSG00000272070* as a conserved, aging-associated lncRNA that modulates microglial inflammatory states through interaction with PU.1, thereby linking glial lncRNA regulation to mechanisms of neurodegeneration.

## Results

### lncRNA 3222401L13Rik regulates cytokine production and phagocytic activity of mouse microglia

In a recent study we used young and aged mice to identify hippocampal glia-specific lncRNAs that have human homologs and show altered expression during aging. Our top candidate was *3222401L13Rik* (**Fig. 1A**), a previously unstudied lncRNA that was elevated in glial cells of the aged hippocampus {Schröder, 2025}. This increase was mainly due to astrocytes, and functionally we found that 3222401L13Rik controls gene expression through its interaction with the transcription factor Neuronal PAS Domain Protein 3 (Npas3). Moreover, we found that higher levels of *3222401L13Rik* act as a compensatory mechanism to support neurons and synapses {Schröder, 2025}. However, strong expression of *3222401L13Rik* was also observed in microglia, and in fact, the baseline expression of *3222401L13Rik* in microglia and astrocytes was similar {Schröder, 2025}. Apart from our recent work on astrocytes {Schröder, 2025}, nothing is known about its role in microglia. Therefore, we set out experiments to investigate the function of 3222401L13Rik in these cells.

**Fig. 1.**
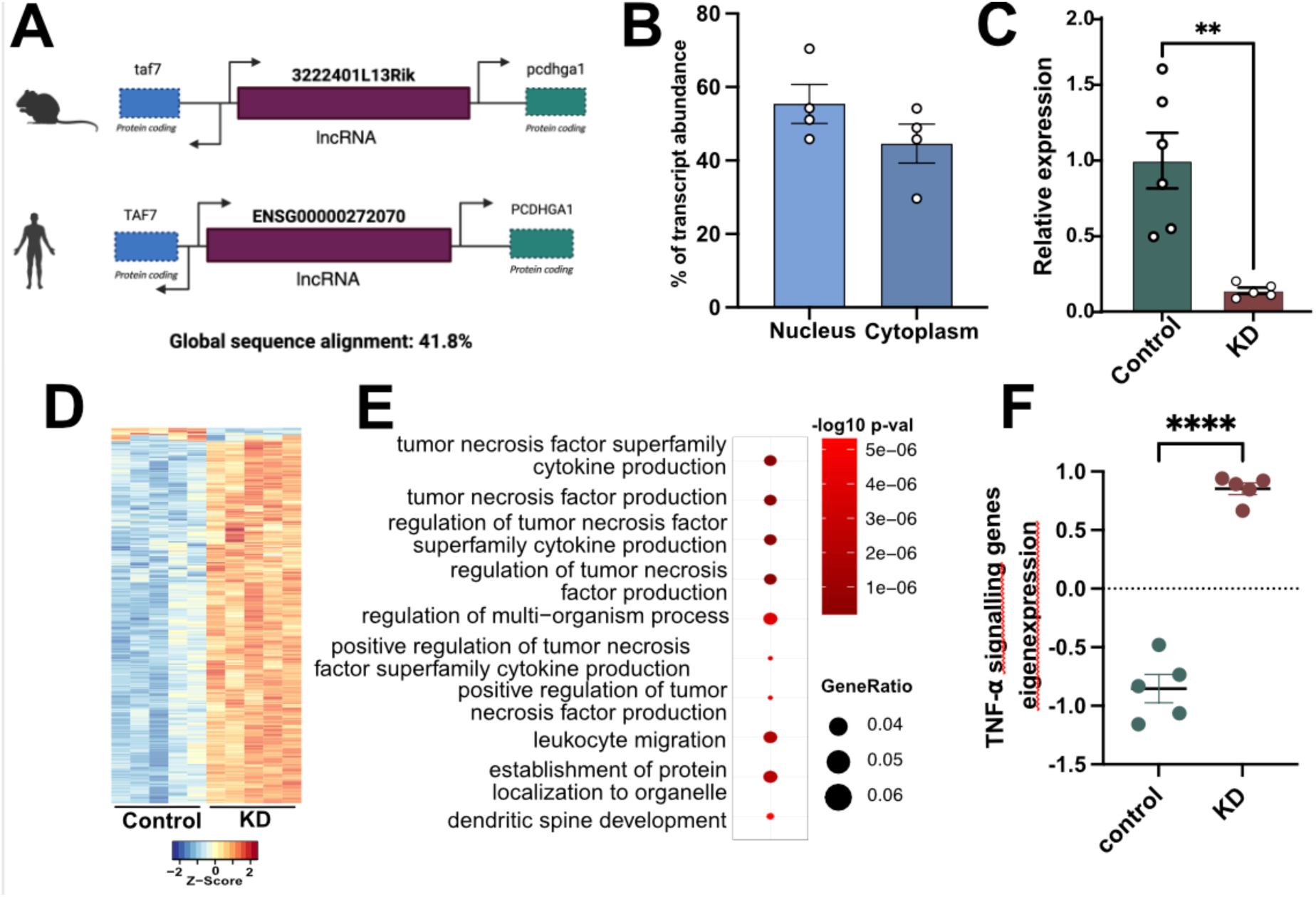
lncRNA 3222401L13Rik regulates genes related to TNF signaling in microglia. **A.**Schematic representation of the genomic localization of 3222401L13Rik in the mouse and human genome. **B.** Bar plots showing the expression of 3222401L13Rik in nuclear and cytoplasmic fractions of primary microglia (n=4). **C.** RT-qPCR quantification of 3222401L13Rik expression after ASO–mediated knockdown (KD) in primary microglia ( n=6 controls & n=5 KD; **p < 0.01, unpaired tTest). **D.** Heatmap of differentially expressed genes from total RNA sequencing after ASO - mediated KD of 3222401L13Rik in primary microglia. **E.** GO terms that are enriched among upregulated genes following 3222401L13Rik KD in primary microglia. **F.** Eigenexpression of genes associated with Tnfα signalling related GO terms from the total RNA sequencing in control and 3222401L13Rik KD primary microglia (n = 5, ****p < 0.0001, unpaired tTest). Error bars indicate SEM.

lncRNAs perform their regulatory functions depending on their compartment-specific localization within the cell {Statello, 2020}. To investigate the function of 3222401L13Rik in microglia, we first analyzed its subcellular localization by performing qPCR on nuclear and cytoplasmic fractions of microglia. The analysis showed that 3222401L13Rik was detectable in both the cytoplasm and the nucleus, with slightly higher expression in the nucleus (**Fig. 1B**). This nuclear localization suggests a role in gene expression control, similar to what we previously observed in astrocytes {Schröder, 2025}.

To test whether 3222401L13Rik regulates transcription in the nucleus, we generated antisense oligonucleotides (ASOs), more specifically, Gapmer ASOs that accumulate and act more effectively in the nucleus than other RNA-targeting oligonucleotides {Monia, 1992} {Crokke, 2021}, to knock down (KD) 3222401L13Rik in primary microglia. We used one ASO targeting 3222401L13Rik and a scrambled control ASO with no RNA target. qPCR confirmed that 3222401L13Rik levels were efficiently reduced in primary microglia (**Fig. 1C**).

Next, to investigate transcriptomic changes after KD of *3222401L13Rik*, we performed total RNA sequencing in primary microglia. Differential gene expression analysis revealed widespread transcriptomic alterations. Notably, we observed almost exclusively upregulated transcripts, with 384 genes upregulated and only 12 genes downregulated after *3222401L13Rik* KD (**Fig. 1D, Supplementary Table 1)**. Gene Ontology (GO) analysis showed that these upregulated genes were mainly associated with processes related to tumor necrosis factor (TNF) signaling and cytokine production, including “*regulation of TNF production*”, “*TNF superfamily cytokine production*” and “*positive regulation of TNF signaling*” (**Fig. 1E, Supplementary Table 2)**. These data suggest that *3222401L13Rik* plays an important role in TNF signaling. In line with this, we observed a strong increase in genes associated with TNFα signaling **(Supplementary Table 3)**, the principal form of TNF expressed in microglia {Gao, 2023}, when their combined expression was calculated as eigenexpression (**Fig. 1F**).

In addition, enriched terms such as “l*eukocyte migration*” and “*dendritic spine developmen*t” suggest that loss of *3222401L13Rik* not only enhances microglial inflammatory pathways but may also influence immune–neuron interactions and synaptic regulation.

We confirmed the increased expression of Tnf⍺ via qPCR (**Fig. 2A**). An increase in *Tnf⍺* expression indicates enhanced neuroinflammatory signaling, reflecting a shift of microglia toward a pro-inflammatory state {Chen, 2019} {Neniskyte, 2014}. In addition, TNF⍺ is closely linked to microglial phagocytosis, suggesting that decreased *3222401L13Rik* levels may influence the clearance of cellular debris-processes highly relevant in aging and neurodegeneration. Therefore, we performed a phagocytic activity assay comparing microglia after *3222401L13Rik* KD to control cells. We observed that KD of *3222401L13Rik* in microglia increased their phagocytic activity (**Fig. 2B**).

**Figure 2.**
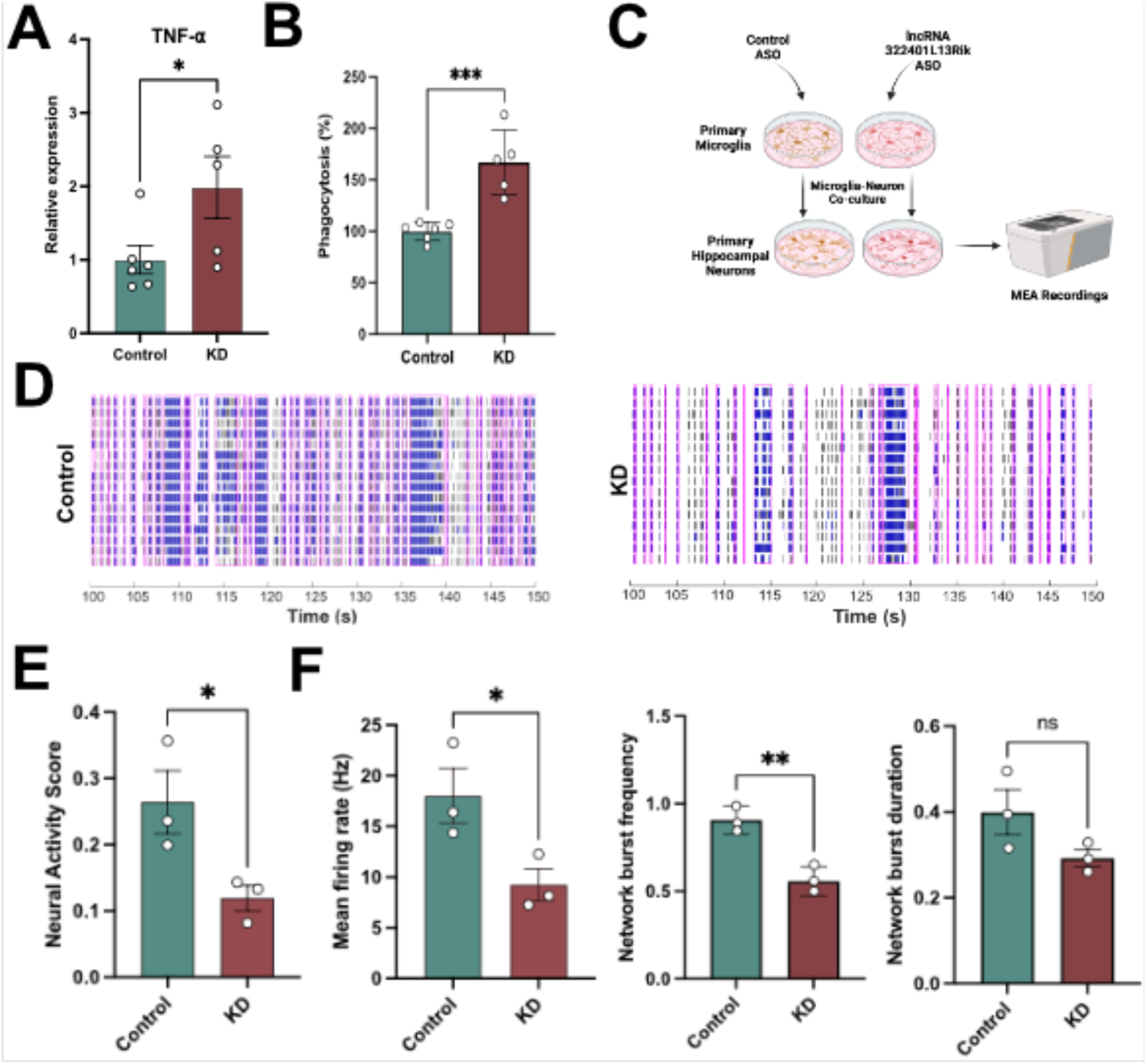
lncRNA 3222401L13Rik regulates cytokine production and phagocytic activity in mouse microglia. **A.** RT-qPCR quantification of Tnfα in primary microglia after 3222401L13Rik knockdown (KD) using ASOs (n = 6 controls & n= 5 KD; *p < 0.05, unpaired t-test). **B.** Bar plot showing phagocytosis (%) in primary microglia after 3222401L13Rik KD (n = 6 controls & n = 5 KD; ***p < 0.001, unpaired t-test). **C.** Schematic overview of the experimental design for the microglia–neuron co-culture experiment. **D.** Representative raster plots showing network activity of microglia–neuron co-cultures with microglia in which ASOs were used to knock down 3222401L13Rik (KD) and the corresponding control cultures. **E.** Neural activity score calculated from MEA recordings (n=3). **F.** Bar charts quantifying mean firing rate (left), network burst frequency (middle), and network burst duration (right) from MEA recordings of microglia–neuron co-cultures after 3222401L13Rik KD. (n = 3; **p < 0.01; *p < 0.05; ns, not significant, unpaired tTest).

In the brain, microglia reside in a multicellular environment where they execute immune functions and support neuronal homeostasis. To assess whether microglia-expressed *3222401L13Rik* KD affects neuronal network activity, we established microglia–neuron co-cultures {Kaurani, 2024} and performed microelectrode array (MEA) recordings (**Fig. 2C**). Neurons were cultured on MEA plates and overlaid with primary microglia transfected with either a control ASO or an ASO targeting *3222401L13Rik*. Microglia were added at a density corresponding to 2% of the neuron count. Co-cultures containing *3222401L13Rik*-KD microglia showed altered firing patterns (**Fig. 2D**) and a significantly reduced neuronal activity score, an objective composite metric for MEA recordings {Passaro, 2021} {Castro-Hernandez, 2023}, when compared with controls (**Fig. 2E**). Detailed analysis revealed decreased mean firing rate and network burst frequency, whereas network burst duration remained unchanged (**Fig. 2F**).

Since pro-inflammatory microglia have been implicated in pathological synaptic pruning {Andoh, 2021}, we also analyzed dendritic spine and synapse numbers in our co-culture system. The results revealed that dendritic spine density and synapse density were unaffected in primary hippocampal neurons co-cultured with 3222401L13Rik KD microglia **(Supplementary** Fig. 1). Together, these findings suggest that loss of 3222401L13Rik biases microglia toward a pro-inflammatory, phagocytosis-competent state that depresses network drive through functional mechanisms such as cytokine-mediated modulation of synaptic efficacy and/or selective clearance of presynaptic or perisynaptic elements, rather than through structural synapse loss.

### lncRNA 3222401L13Rik human homolog ENSG00000272070 regulates cytokine production and phagocytic activity of iPSC-derived human microglia-like cells

To test whether the human homolog of *3222401L13Rik* (*ENSG00000272070*) has similar effects, we used iPSC-derived human microglia-like cells (iMGLs), a well-established human microglia model {Kaurani, 2023}. We first analyzed the subcellular localization of *ENSG00000272070* in iMGLs. QPCR of nuclear and cytoplasmic fractions showed that, similar to mouse microglia, ENSG00000272070 localizes to both compartments, with slightly higher levels in the nucleus (Fig. 3A). Next, we generated antisense oligonucleotides (ASOs) to knock down (KD) *ENSG00000272070* in iMGLs (Fig. 3B) and measured TNFα by qPCR. Consistent with the mouse data (see Fig. 2A), *TNFα* levels were increased in iMGLs upon *ENSG00000272070* KD (Fig. 3C). In addition, *ENSG00000272070* KD increased phagocytic activity in iMGLs, in agreement with the findings in murine cells (Fig. 3D).

**Figure 3.**
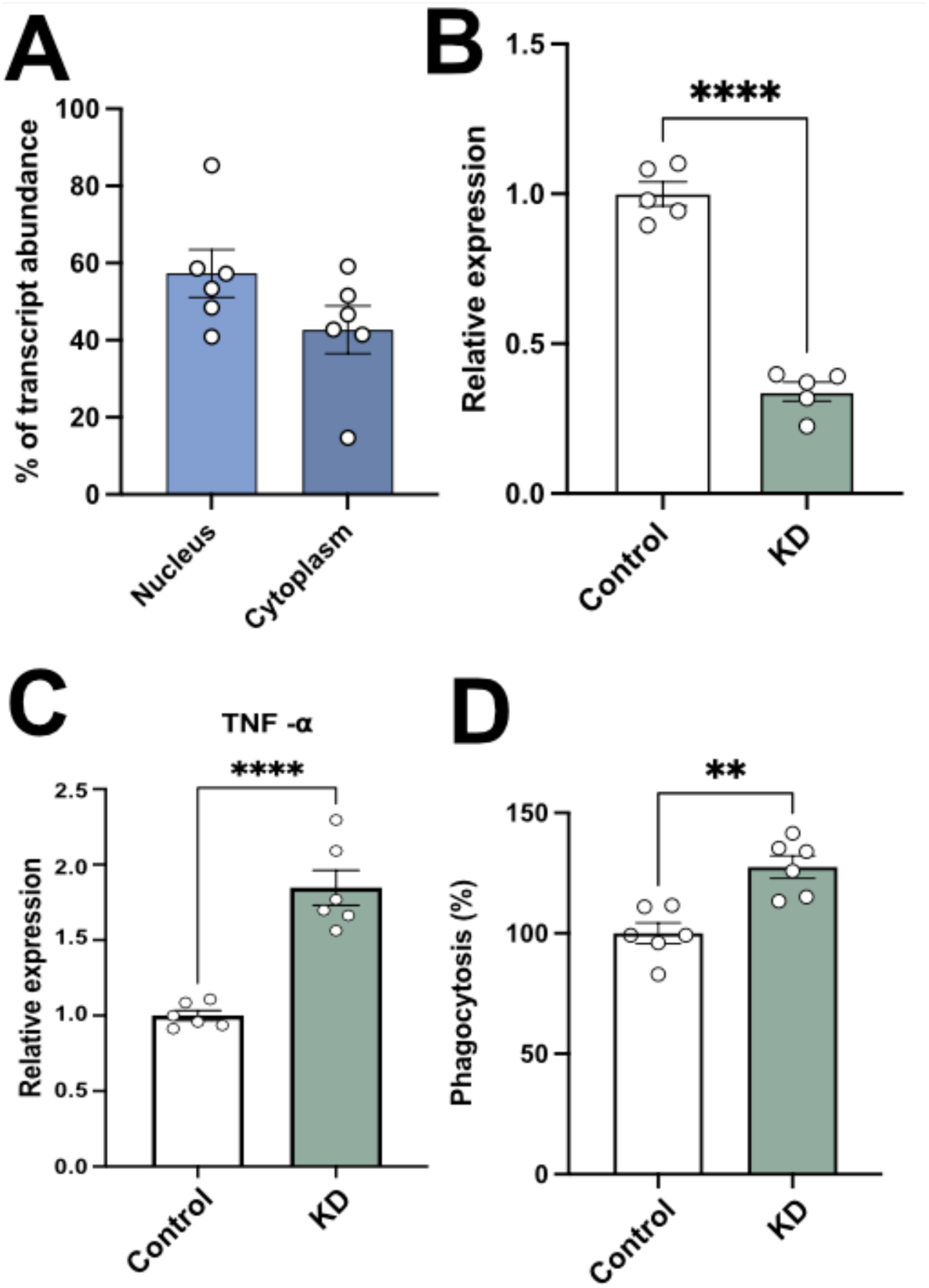
ENSG00000272070 (human homolog of lncRNA 3222401L13Rik) regulates cytokine production and phagocytosis in iMGLs. **A.** Expression of ENSG00000272070 in nuclear and cytoplasmic fractions of iMGLs (n = 6). **B.** qPCR quantification of ENSG00000272070 after ASO-mediated knockdown (KD) in iMGLs (n = 5; ****p < 0.0001; unpaired t-test). **C.** qPCR quantification of TNFα in iMGLs after ENSG00000272070 KD (n = 6; ****p < 0.0001; unpaired t-test). **D.** Bar plot showing phagocytosis (%) by iMGLs after ENSG00000272070 KD (n = 6; **p < 0.01; unpaired t-test).

These findings suggest that the human lncRNA *ENSG00000272070* and its mouse homolog *3222401L13Rik* exhibit, at least in part, similar functions, namely, the regulation of *TNFα* signaling and phagocytosis.

### lncRNA 3222401L13Rik/ENSG00000272070 is deregulated in AD and induces transcriptomic and functional changes similar to AD

Next we set out experiments to analyze the potential implication of the human homologue of *3222401L13Rik, ENSG00000272070,* in AD. First, we quantified its expression by qPCR in the postmortem prefrontal cortex (Brodmann area 9, BA9) from AD cases (Braak stage IV) and age-matched controls. ENSG00000272070 levels were significantly decreased in AD (Fig. 4A). We also examined other neurological disorders and found no significant differences in brains from individuals with Parkinson’s disease, frontotemporal dementia, or schizophrenia (Supplementary Fig. 2).

**Figure 4.**
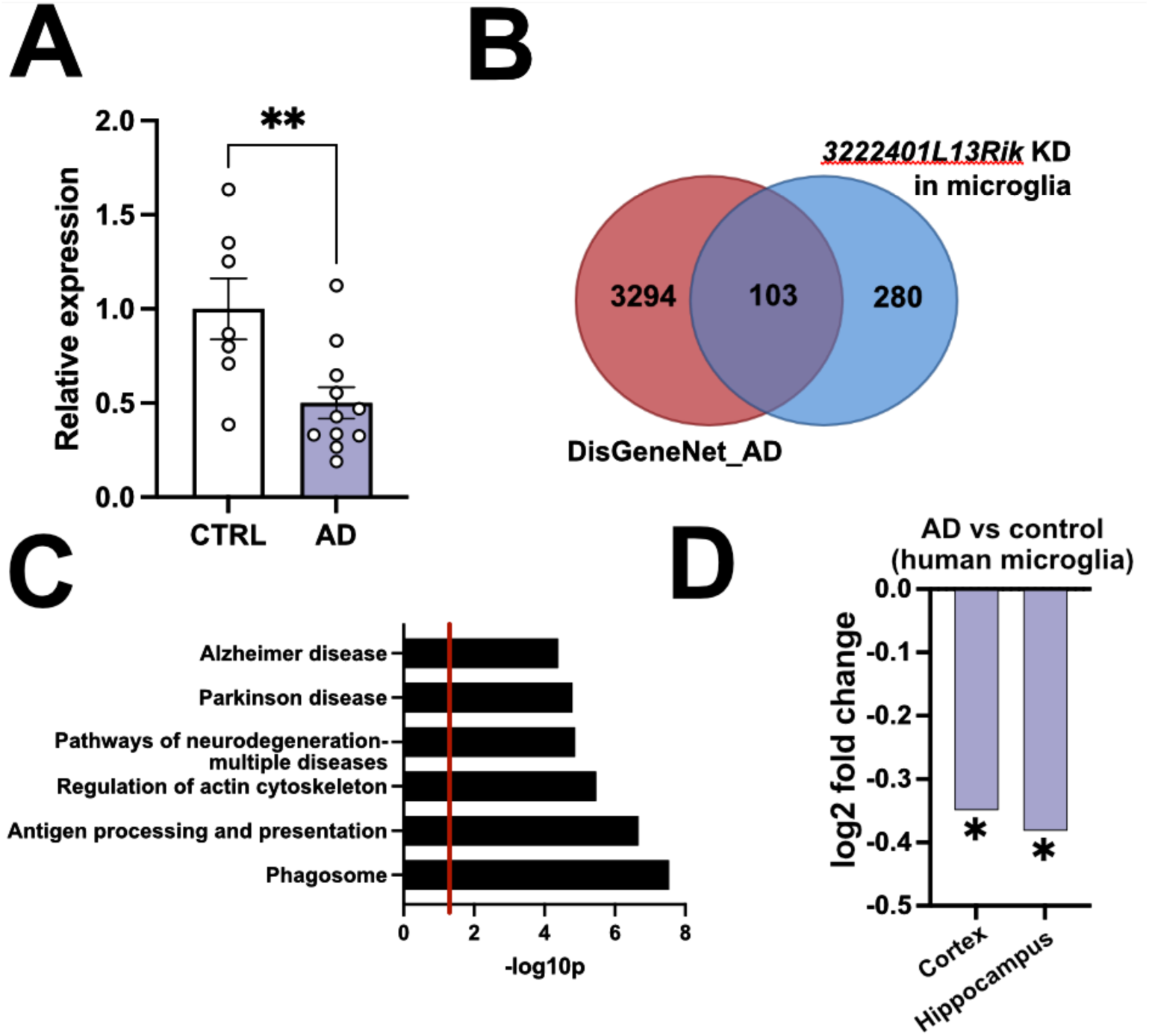
Loss of lncRNA 3222401L13Rik in microglia induces transcriptomic changes similar to AD. **A.** Bar plot showing ENSG00000272070 expression in postmortem prefrontal cortex (BA9) from controls (n = 7) and AD cases (n = 11). **p < 0.01; unpaired tTest**. B.** Venn diagram showing the overlap between AD-associated genes from DisGeNET (AD subset) and genes upregulated in primary microglia after 3222401L13Rik knockdown (KD). **C.** Gene Ontology (GO) terms associated with the 103 shared genes between 3222401L13Rik KD in primary mouse microglia (PMMs) and DisGeNET-AD genes. **D.** Bar chart showing the log2 fold change of ENSG00000272070 in microglia, when comparing human brain snucRNAseq data from AD vs control individuals. (*P < 0,05; Likelihood Ratio Test). The snucRNAseq dataset is from {Jiang, 2020}.

Given the important roles of microglia in AD, we next asked whether the gene expression changes observed in primary microglia after 3222401L13Rik KD (see Fig. 1) overlap with those reported in AD. We obtained AD-associated genes from DisGeNET (AD subset; an integrative resource that aggregates gene–disease associations from curated databases, Genome-wide association studie (GWAS), animal models, and literature mining) **(Supplementary Table 4)** {Piñero, 2017}. We observed that 103 of the 303 genes upregulated after 3222401L13Rik KD in primary microglia are associated with AD according to DisGeNET (Fig. 4B**, Supplementary Table 5)**. GO term analysis of these genes revealed enrichment for “*Neurodegeneration*”, “*Alzheimer’s disease*”, and “*Phagosome*” pathways (Fig. 4C**, Supplementary Table 6)**.

Finally, we analyzed the expression of *ENSG00000272070* in a single-cell/nucleus RNA-seq database {Jiang, 2020} and found that *ENSG00000272070* was significantly decreased in microglial cells of the human hippocampus and cortex when comparing AD patients to controls (Fig. 4D**, Supplementary Table 7)**.

### lncRNA 3222401L13Rik interacts with microglia-specific transcription factor PU.1

Our findings suggest that lncRNA *3222401L13Rik* modulates microglial function by orchestrating gene-expression programs. Long noncoding RNAs can regulate gene expression through several mechanisms, including acting as molecular decoys or recruiters for transcription factors {Statello, 2021} {Schröder, 2024} {Pradhan, 2025}. To test this, we performed an enrichment analysis of genes upregulated after *3222401L13Rik* knockdown in primary microglia and identified *Spi-1 proto-oncogene* (*Pu.1/Spi1*) as the most significantly enriched transcription factor (Fig. 5A). *Pu.1* is a myeloid-lineage transcription factor essential for microglial identity and function {Cakir, 2022} {Smith, 2013} and has been genetically and functionally linked to AD, with higher *PU.1* levels being associated with an increased AD risk, while lower levels are protective {Huang, 2017} {Pimenova, 2020} {Ralvenius, 2024}. In line with this, *PU.1* expression was found to be elevated in human AD brains, which coincided with increased expression of PU.1 target genes in human microglia {Rustenhoven, 2018a}. We confirmed higher *PU.1* mRNA levels in human AD brains using the AGORA dataset, revealing significant increases in posterior cingulate cortex (PCC), dorsolateral prefrontal cortex (DLPFC), and parahippocampal gyrus (PHG) (Fig. 5B). To test for a physical association between *3222401L13Rik* and PU.1, we performed RNA immunoprecipitation (RIP) in primary microglia using PU.1 antibodies, with TATA-Binding Protein-Associated Factor (TAF) and IgG as controls. *3222401L13Rik* was significantly enriched in the PU.1 IP but not in control IPs (Fig. 5C). RIP in human iMGLs similarly showed that *ENSG00000272070* binds PU.1 (Fig. 5D). These data suggest that *3222401L13Rik* and *ENSG00000272070* regulate PU.1-dependent gene expression, likely by acting as molecular decoys for PU.1, since loss of *3222401L13Rik* led almost exclusively to increased gene expression in microglia (see Fig. 1). To test this further, we re-analyzed a previously published dataset in which the authors performed PU.1 overexpression and knockdown (KD) in BV2 microglia {Pimenova, 2020}.

**Figure 5.**
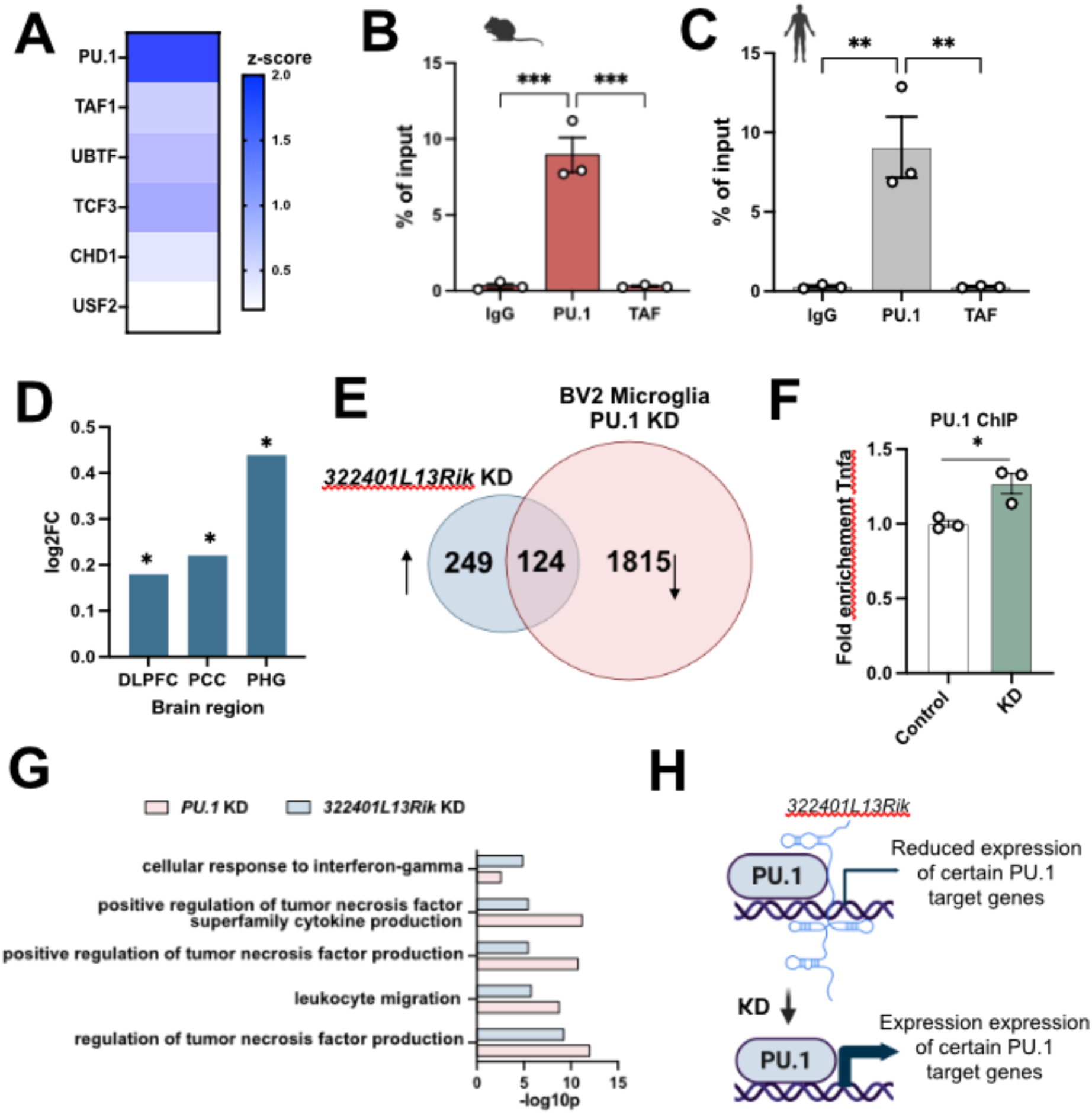
lncRNA *3222401L13Rik* interacts with the PU.1 transcription factor. **A.** Enrichment analysis for transcription factors (TF) based on common upregulated genes upon lncRNA KD in primary microglia. Data were obtained from the EnrichR dataset. **B.** Bar chart showing the quantification of RNA immunoprecipitation (RIP) for *3222401L13K* in primary microglia probing for *PU.1, TAF* and IgG control. (n= 3; *\*\*\*p < 0.001; one way ANOVA)*. **C.** Bar chart showing the quantification of RNA immunoprecipitation (RIP) for *3222401L13K* in human iPSC-derived microglia probing for *PU.1, TAF* and IgG control. (n = 3; *\*\*p < 0.01; one way ANOVA).* **D.** Bar plot showing the differential expression of *PU.1* TF in postmortem AD brain across different brain areas. Data was obtained from the AGORA AD database. *\*p < 0.05*. **E.** Overlap showing the commonly deregulated genes between *3222401L13Rik* KD in primary microglia and *PU.1* KD in BV2 microglia from {Pimenova, 2020}. **F.** ChIP for PU.1 TF in primary microglia treated with negative control gapmer (control) or *3222401L13Rik* gapmer (KD) followed by qPCR for TNFa. (n = 3; *\*p < 0.05; unpaired tTest)*. **G.** Common GO terms for sderegulated genes in *PU.1* KD and *3222401L13Rik* KD microglia. **H.** Schematic depiction of the predicted mode of function of *3222401L13Rik* in microglia.

In agreement with the original findings, PU.1 overexpression had only minor effects on gene expression, with just 30 differentially expressed genes (FDR < 0.05, log₂FC ≥ 0.5) **(Supplemental** Fig. 3A**; Supplemental Table 8)**. In contrast, *PU.1* KD led to substantial gene-expression changes **(Supplemental** Fig. 3B**; Supplemental Table 9)**.

We reasoned that genes downregulated upon *Pu.1* KD (1,939 genes) and upregulated upon *3222401L13Rik* KD (373 genes) may represent PU.1-dependent genes that are co-regulated via *3222401L13Rik*.

Cross-referencing our RNA-seq dataset from *3222401L13Rik* KD with genes downregulated upon *Pu.1* KD in BV2 microglia revealed that approximately 33% (124 genes) of the *3222401L13Rik*-regulated genes overlap with PU.1 targets, including *Tnf⍺* (Fig. 5E**; Supplementary Table 10)**. To confirm this, we treated primary microglia with *3222401L13Rik* ASOs or control oligos and performed PU.1 chromatin immunoprecipitation (ChIP) followed by qPCR for *Tnf⍺*. The data show increased PU.1 levels at the *Tnf⍺* promoter upon *3222401L13Rik* KD (Fig. 5F).

Moreover, when we subjected the 124 genes potentially co-regulated by PU.1 and *3222401L13Rik* to GO term analysis and compared the identified signaling pathways to those affected by *3222401L13Rik* KD, key shared pathways linked to TNF⍺ signaling, such as “*regulation of tumor necrosis factor production*” and “*positive regulation of tumor necrosis factor superfamily cytokine production*”, were identified (Fig. 5G**; Supplementary Table 11**).

Together, these data support a model in which 3222401L13Rik acts as a decoy for PU.1, restraining its activity at selected gene, especially those linked to TNF⍺ signaling, while loss of 3222401L13Rik, as observed in AD patients, enhances the expression of these genes (Fig. 5H).

## Discussion

We identify *3222401L13Rik* as a glia-enriched lncRNA that regulates microglial state. Loss of *3222401L13Rik* in primary mouse microglia reprograms transcription toward pro-inflammatory cytokine pathways and increases phagocytic activity, accompanied by *Tnfα* upregulation and reduced neuronal firing in microglia–neuron co-cultures without overt synapse loss. We further nominate *ENSG00000272070* as a conserved human counterpart and its knockdown in iPSC-derived microglia also elicits increased phagocytosis and elevated *TNFα* levels. Moreover, *ENSG00000272070* expression is reduced in the human AD brain. Mechanistically, we show that *3222401L13Rik* interacts with PU.1, an AD-associated myeloid transcription factor {Huang, 2017} {Pimenova, 2020} {Ralvenius, 2024}, and that *3222401L13Rik*-dependent genes substantially overlap PU.1 targets. Collectively, these data support a novel lncRNA–PU.1 axis that tunes microglial activation during aging and disease in mice and humans.

### Cell-type–specific modes of action of 3222401L13Rik

To date, only one other study has addressed the role of *3222401L13Rik*. Specifically, our group recently identified *3222401L13Rik* as a glia-enriched lncRNA that is upregulated in the aging mouse hippocampus {Schröder, 2025}. While *3222401L13Rik* is not expressed in neurons, it is detectable in astrocytes, microglia, and oligodendrocytes. In astrocytes, we showed that *3222401L13Rik* regulates neuronal support and synaptic organization through interaction with the transcription factor Npas3 {Schröder, 2025}. Consistent with this, our present data demonstrate that in microglia, *3222401L13Rik* also contributes to gene regulation via interaction with a transcription factor PU.1. Although this does not exclude the likely possibility that *3222401L13Rik* influences microglia function through additional mechanisms, this mode of action is consistent with other lncRNAs known to modulate gene expression through interactions with transcription factors {Marchese, 2017} {Statello, 2021}.

The finding that *3222401L13Rik* interacts with different transcription factors in astrocytes and microglia aligns with our observation that *3222401L13Rik* KD produces distinct transcriptional profiles in these cell types. KD of *3222401L13Rik* in microglia led almost exclusively to upregulated genes, whereas KD in astrocytes predominantly resulted in downregulated genes {Schröder, 2025}. A direct comparison revealed minimal overlap between the two datasets. Among 321 genes significantly upregulated in astrocytes {Schröder, 2025} and 385 genes significantly upregulated in microglia, only five genes were shared **(Supplementary Table 10)**. This difference may partly reflect the fact that PU.1 is not expressed in astrocytes {Cahoy, 2008}. Further studies are required to elucidate the precise mechanisms by which *3222401L13Rik* orchestrates distinct transcriptional programs in astrocytes and microglia. It will also be important to determine the structural organization of *3222401L13Rik* and to identify the specific regions mediating interactions with PU.1 or Npas3. Moreover, RNA molecules can adopt different structural conformations depending on the cellular environment {Wan, 2014}. Thus, mapping the structure of *3222401L13Rik* in astrocytes and microglia may also help to reveal if cell type-specific RNA folding might contribute to its divergent functions. However, these experiments go beyond the scope of this study and will be addressed in future research.

### 3222401L13Rik modulates inflammatory microglial states

Our finding that *3222401L13Rik* regulates microglial function aligns with prior reports implicating other lncRNAs in microglia biology. For example, *Neat1* is strongly upregulated by cellular stress and inflammatory cues in microglia and can drive pro-inflammatory programs, including increased TNF-α, IL-1β, and IL-6 expression {Pan, 2022}. Consistent with this axis, we observed that reducing *3222401L13Rik* elevated expression of genes linked to pro-inflammatory processes, with TNF-α signaling emerging as a central pathway. This suggests that, under physiological conditions, *3222401L13Rik* may act as a molecular counterplayer to *Neat1* and other pro-inflammatory drivers operating through, at least in part, TNF-α signaling. Notably, microglia rapidly release TNF-α in response to danger cues (e.g., LPS, Aβ, injury), influencing NLRP3 inflammasome activation, NF-κB signaling, and phagocytosis {Gao,2023}, the latter of which is also increased after *3222401L13Rik* KD in murine micoglia and upon *ENSG00000272070* KD in human iPSC-derived microglia. Together, these observations support a model in which *3222401L13Rik* and *ENSG00000272070* function as a molecular decoy that fine-tunes and restrains pro-inflammatory microglial states, at least in part by limiting PU.1 occupancy at the TNF-α promoter, as shown by our data. This is in agreement with data showing that elevated TNF-α is a hallmark of several neurodegenerative diseases, including AD, and can promote chronic neuroinflammation and synaptic dysfunction, when unchecked {Heneka, 2015} {Heppner, 2015} and numerous data showing that PU.1 drives, amongst other genes, Tnf-α expression in the context of neurodegenerative diseases {Ralvenius, 2024} {Xu, 2025}. This view aligns also with our observation that *3222401L13Rik* KD in microglia suppresses neuronal network activity without reducing spine numbers, as pro-inflammatory TNF-α signaling has been shown to influence homeostatic synaptic plasticity, including lower network plasticity without immediate spine pruning {Tancredi, 1992} {Butler, 2004}. That said, the effects of TNF-α on neuronal plasticity are complex and context-dependent, and TNF-α has also been reported to increase neuronal excitability {Stellwagen, 2006}. Nevertheless, the literature supports the idea that TNF-α released by microglia and also astrocytes fine-tunes neuronal function {Stellwagen, 2006}, suggesting that *3222401L13Rik* may act as a homeostatic brake that normally helps to stabilize circuits but, when deregulated in AD, contributes to pushing networks into maladaptive states. At present this remains speculative, and further work will be necessary to define the precise mechanisms by which *3222401L13Rik* and *ENSG00000272070* contribute to the observed phenotypes. It is also important to note that our current data derive from acute manipulations, whereas AD is characterized by chronic neuroinflammatory dynamics. Thus, longitudinal and *in vivo* studies will be essential to further establish causal relevance and temporal progression.

### A 3222401L13Rik –PU.1 axis shaping inflammatory gene programs

When analyzing the genes deregulated in microglia upon *3222401L13Rik* KD, PU.1 emerged as the most significant transcription factor likely to explain the regulation of these genes. In agreement with this, we found that *3222401L13Rik* interacts with PU.1, and that *3222401L13Rik* KD in microglia increased the binding of PU.1 to the *Tnf-α* promoter, which correlated with elevated *Tnf-α* expression. GWAS have identified the *PU.1* gene locus as being associated with AD risk and a specific allelic variant that leads to lower *PU.1* expression is associated with a delayed onset of AD {Huang, 2017}. These data provide strong evidence that a therapeutic approach aimed at reducing PU.1 in microglia could be a suitable strategy to mitigate neuroinflammatory processes involved in AD pathogenesis.

In line with this, reducing PU.1 levels has been shown to decrease the microglial inflammatory response {Pimenova, 2020} {Rustenhoven, 2018} {Ralvenius, 2023}. Taken together, these observations indicate that a key function of *3222401L13Rik* in microglia is to restrict PU.1 activity by preventing it from binding to its target genes. Although we experimentally demonstrate this for only one target gene, namely *Tnf-α*, a comparison of microglial gene expression following *3222401L13Rik* KD with that after *PU.1* KD in BV2 microglia {Pimenova, 2020} revealed that a significant proportion of the genes regulated by *3222401L13Rik* are also PU.1 targets, providing further support for the view that *3222401L13Rik* acts as a molecular decoy for PU.1 in microglia.

In summary, our study identifies the aging-related lncRNA *3222401L13Rik* as a regulator of microglial activation through interaction with the AD-associated transcription factor PU.1. The human homolog *ENSG00000272070* is reduced in AD brains, and gene-expression changes following its knockdown parallel AD-associated transcriptomic shifts. Together, these findings reveal a lncRNA-mediated mechanism of PU.1 regulation that shapes microglial inflammatory states and highlight *3222401L13Rik*/*ENSG00000272070* as potential targets for microglia-oriented therapeutic strategies in Alzheimer’s disease.

## Methods

### Animal experiments

Animals were (Janvier Labs, Le Genest St Isle, France) were kept under a 12h/12h light/dark cycle, in standard single cages with food and water provided ad libitum. The protocol to generate primary neurons and to collect tissue from mice was approved by the Lower Saxony State Office for Consumer Protection and Food Safety (AZ 22.00203).

### Isolation of nuclei from mouse and human brain tissue

The isolation of nuclei from mouse and human brain tissue was performed according to our previous publication {Michurina, 2022} with minor modifications. Briefly, tissue was homogenized using a Dounce homogenizer in 500 µl EZ Prep lysis buffer (Sigma) with 30 strokes (for mouse tissue). The homogenate was transferred into a 2 ml Eppendorf tube, and the volume was adjusted to 2 ml with lysis buffer, followed by incubation on ice for 7 min. The homogenate was centrifuged for 5 min at 500 × g at 4 °C, and the supernatant was discarded. The nuclei pellet was resuspended in 2 ml lysis buffer and incubated on ice for 5 min, followed by another centrifugation step (5 min, 500 × g, 4 °C). The resulting pellet was resuspended in 1.5 ml nuclei storage buffer (NSB; 1× PBS, 0.5% BSA, 1:200 RNase inhibitor, 1:100 Roche protease inhibitor) and centrifuged again for 5 min (500 × g, 4 °C). The obtained pellet was resuspended in 1 ml NSB and stained with anti-NeuN-AlexaFluor® for 1 h at 4 °C, followed by centrifugation for 5 min (500 × g, 4 °C). The pellet was washed once with 500 µl NSB, resuspended in 300–500 µl NSB depending on pellet size, and stained with 7-AAD (1:100). Samples were passed through a 40 µm filter into FACS tubes and gently vortexed before sorting. Sorting of NeuN-positive and NeuN-negative nuclei was performed on a FACSAria III (BD Biosciences). Sorted nuclei were counted using a Countess II FL Automated Cell Counter. Finally, sorted nuclei were centrifuged and resuspended in TRIzol-LS for RNA isolation for qPCR or RNA sequencing.

### RNA sequencing and analysis

Libraries were prepared using the SMARTer Stranded Total RNA-seq v2 – Pico Input Kit (Takara Bio) with 8 ng of input RNA. Libraries were amplified for 13 cycles, and library size and quality were assessed using a Bioanalyzer (Agilent Technologies). Multiplexed libraries were sequenced on an Illumina NextSeq 2000 platform generating 50 bp single-end reads. Sequencing data were processed using a customized in-house pipeline. bcl2fastq (v2.20.2, Illumina) was used for adapter trimming and conversion of per-cycle BCL files to per-read FASTQ files. Quality control of raw reads was performed using FastQC (v0.11.5) (http://www.bioinformatics.babraham.ac.uk/projects/fastqc/). Reads were aligned to the mouse reference genome (mm10) using STAR (v2.5.2b), and read counts were generated with featureCounts (v1.5.1). Differential gene expression analysis was performed using DESeq2 (v1.30.0) {Love et al., 2014}, incorporating RUVSeq to correct for unwanted variation. Gene Ontology (GO) term enrichment was conducted using the ClueGO plugin in Cytoscape, and transcription factor enrichment analysis was carried out with Enrichr (https://maayanlab.cloud/Enrichr/).

### Primary microglia culture

Primary mouse microglia were prepared as previously described {Lian et al., 2016} with minor modifications. Briefly, CD1 pups aged postnatal day 1–2 were decapitated, and brains were rapidly removed. The meninges were carefully stripped away in dissection medium containing HBSS, 1% 1 M HEPES, 3 g/L glucose, and 1% penicillin/streptomycin, and the forebrains (cortex and hippocampus) were dissected. Three half-forebrains (i.e., tissue from 1.5 pups) were transferred into 50 ml Falcon tubes containing 5 ml of dissection medium. The tissue was gently dissociated using a 1 ml pipette and digested for 15 min at 37 °C in 2.5% trypsin, followed by DNase I treatment. Cells from each Falcon tube were plated into poly-D-lysine (PDL)-coated T75 flasks and cultured in basal medium (DMEM supplemented with 10% fetal bovine serum and 1% penicillin/streptomycin) in a humidified incubator at 37 °C with 5% CO₂ for 7–10 days. To harvest microglia, flasks were placed on an Incu-Shaker Mini (Benchmark) and shaken at 160 rpm for 3 h to detach microglia growing on top of the astrocytic monolayer. The supernatant containing detached microglia was collected, centrifuged at 400 × g for 5 min, and the pellet was resuspended in basal medium. Microglia were plated at a density of 50,000 cells/cm² on culture dishes, and the medium was completely replaced after 2 h to remove unattached cells. Cultures were maintained under standard incubator conditions. To obtain additional microglia, the medium in the T75 flasks was replaced with fresh medium, and cells were harvested again after 72 h using the same shaking procedure.

### Primary neuron culture

Primary hippocampal neurons (PHNs) were prepared from CD1 embryos at embryonic day 17 (E17). Briefly, pregnant CD1 females were euthanized by pentobarbital overdose, and embryonic brains were rapidly isolated. The meninges were carefully removed, and hippocampi were dissected under a stereomicroscope. To obtain a single-cell suspension, the tissue was digested with trypsin, followed by DNase I treatment. Cells were plated at a density of 60,000 cells/cm² on poly-D-lysine–coated multi-well plates and maintained in Neurobasal medium supplemented with 1× B27 and 1 mM GlutaMAX. PHNs were used for experiments at days in vitro (DIV) 10–12.

### Human derived iPSC microglia like cells culture

iPSC-derived human microglia-like cells (iMGLs) were differentiated as previously described {Kaurani, 2024}. Briefly, 3 × 10⁶ induced pluripotent stem cells (iPSCs) (cell line: KOLF2.1J, Jackson Laboratory) were seeded into AggreWell 800 plates (STEMCELL Technologies) to form embryoid bodies (EBs) and cultured in medium supplemented with 50 ng/ml BMP4 (Miltenyi Biotec), 50 ng/ml VEGF (Miltenyi Biotec), and 20 ng/ml SCF (R&D Systems). On day 4, EBs were transferred to 6-well plates (15 EBs per well) and further differentiated in medium containing 100 ng/ml M-CSF (Miltenyi Biotec), 25 ng/ml IL-3 (Miltenyi Biotec), 2 mM GlutaMAX (Invitrogen), and 0.055 mM β-mercaptoethanol (Thermo Fisher Scientific). Fresh medium was added weekly. Microglial precursors appeared in the supernatant after approximately 4 weeks and were collected using a 40 µm cell strainer. Precursors were plated at a density of 20,000 cells/cm² and differentiated into iMGLs in the presence of 100 ng/ml M-CSF and 25 ng/ml interleukin-34 (IL-34).

### Antisense LNA Gapmers

Knockdown of 3222401L13Rik in mouse microglia and ENSG00000272070 in human iMGLs was performed using antisense LNA gapmers targeting the respective lncRNAs. The lncRNA-specific and negative control (NC) gapmers were designed and synthesized by Qiagen. The sequences used were as follows: NC: GCTCCCTTCAATCCAA 3222401L13Rik: AGCTTGGTCATTTGAT ENSG00000272070: GGACTTCTTCCTCTGT Mouse microglia and iMGLs were transfected with gapmers using Lipofectamine RNAiMAX (Thermo Fisher Scientific) according to the manufacturer’s instructions.

### RNA extraction

Cells were lysed using TRI Reagent (Sigma-Aldrich). For tissue samples, homogenization was performed in TRI Reagent using an Omni Bead Ruptor 24 (OMNI International). Total RNA was isolated with the RNA Clean & Concentrator-5 Kit (Zymo Research) according to the manufacturer’s instructions. RNA concentration and purity were measured using a NanoDrop spectrophotometer (Thermo Fisher Scientific).

### cDNA synthesis and qPCR

cDNA was prepared using the transcription cDNA first strand synthesis kit (Roche) with 10-300 ng RNA. qPCR was performed using LightCycler 480 SYBR Green I Master Mix (Roche) on a LightCycler 480 system (Roche) with primers listed in supplementary table 12. Data was analyzed using the 2-ddCt method.

### Phagocytoses assay

Microglia phagocytoses assay was performed using the Vybrant phagocytosis assay kit (Invitrogen) according to manufacturer’s instructions. The assay was validated in each trial by treating microglia with lipopolysaccharide (LPS) for 4 and 24 hours **(Supplementary** Fig. 4) as a positive control. Phagocytosis was measured using FLUOstar Omega (BMG Biotech).

### Publicly available datasets

For analysis of lncRNA expression publicly available datasets were used. The broad institute single cell ageing mouse brain database {Ximerakis, 2019} was used to analyze lncRNA expression in ageing microglia cells. The RiMOD FTD dataset {Menden, 2023} (https://www.rimod-ftd.org/) was used to analyze lncRNA expression in FTD brains. The AGORA AD database (https://agora.adknowledgeportal.org/) was used to analyze expression of lncRNA in AD brains. The GEO dataset GSE156928 was analyzed to determine expression of lncRNA in PD brains. For identification of PU.1 TF gene targets, we analyzed the PU.1 KD dataset in BV2 microglia {Pimenova, 2020} and single cell data stems from {Jiang, 2020}.

### Nuclear and cytoplasmic fractionation

Nuclear and cytoplasmic fractions from microglia were isolated using the EZ Prep Nuclei Isolation Kit (Sigma-Aldrich) according to the manufacturer’s instructions. Briefly, EZ Lysis Buffer was added to the cells, which were then lysed and collected by thorough scraping. Nuclei were pelleted by centrifugation at 500 × g for 5 min at 4 °C. The nuclei were contained in the pellet, while the supernatant represented the cytoplasmic fraction. TRIzol-LS Reagent (Thermo Fisher Scientific) was added to both fractions, which were then stored at –80 °C until RNA isolation.

### RNA-immunoprecipitation (RNA-IP)

Nuclear and cytoplasmic fractions were obtained from cultured mouse microglia or iMGLs using the EZ Prep Nuclei Isolation Kit (Sigma-Aldrich), as described above. The nuclear fraction was resuspended in TSE buffer (10 mM Tris, 300 mM sucrose, 1 mM EDTA, 0.1% Nonidet P-40, 100 U/ml RNase inhibitor, 1× protease inhibitor), transferred to Bioruptor tubes (Diagenode), and sonicated using a Bioruptor Plus (Diagenode) for five cycles (30 s on / 30 s off). Samples were then incubated on ice for 20 min with occasional vortexing and centrifuged for 10 min at 13,000 rpm at 4 °C. The supernatant was transferred to a fresh tube and flash-frozen at –80 °C. The lysate was precleared with 25 µl Pierce™ Protein A/G Magnetic Beads (Thermo Fisher Scientific) for 1 h at 4 °C to reduce nonspecific binding. Anti-EZH2 antibody (Millipore) or an IgG isotype control (Santa Cruz Biotechnology) was incubated with 50 µl Protein A/G magnetic beads for 2 h at room temperature and washed with RNA-IP buffer (50 mM Tris-HCl, 100 mM NaCl, 32 mM NaF, 0.5% NP-40). Ten percent of the lysate was reserved as input, and the precleared antibody–bead complexes were added to the remaining lysate and incubated overnight (O/N) at 4 °C on a rotator. Beads were then washed five times with RNA-IP buffer and incubated with proteinase K for 1 h at 37 °C. The supernatant was collected, and RNA was extracted using the RNA Clean & Concentrator-5 Kit (Zymo Research).

### Human tissues

For the analysis of lncRNA expression by qPCR, brains (prefrontal cortex, BA9) from control and AD patients were obtained from control (n = 6 females & 1 male; age = 86 ± 4.9 years, PMD = 6.08 ± 2.5 h) and AD patients (n = 10 females & 1 male; age = 77.5 ± 11.5 years, PMD = 13.5 ± 3.88 h; Braak & Braak stage IV) were obtained with ethical approval from the ethics committee of the University Medical Center Göttingen and upon informed consent from the Harvard Brain Tissue Resource Center (Boston, USA).

### Microglia-neuron co-culture

Primary hippocampal neurons were seeded at a density of 50,000 cells/cm² on polyethyleneimine (PEI)-coated MEA plates for electrophysiological recordings or on poly-D-lysine (PDL)-coated 24-well glass-bottom plates (Cellvis) for imaging. Neurons were used for experiments at days in vitro (DIV) 10–12. Primary microglia treated with antisense LNA gapmers were harvested from T75 flasks, and 2,000 microglia were added to each neuronal culture well. After co-culture, plates were maintained under standard incubator conditions (37 °C, 5% CO₂, humidified atmosphere).

### Multielectrode array recordings

To record spontaneous neuronal activity, the Maestro Edge multi-well MEA system (Axion Biosystems) was used. Primary hippocampal neurons were plated on PEI coated wells of 24-well MEA plate as described above. 24h after co-culture with treated microglia (described above) the MEA plate was transferred to the recording chamber maintained at 37°C with 5% CO2. The activity of neurons was recorded for 15 min. using Axion Biosystem’s studio software (setting for frequency cutoff). Data analysis was performed using the Neural metric tool (Axion Biosystems). The neural activity score was calculated as described in {Castro-Hernandez, 2023}.

### Chromatin immunoprecipitation (ChIP)

Primary microglia were cultured in 6-well plates and treated with NC or KD LNPs as described above. To each well, 1 ml of low-sucrose buffer (0.32 M sucrose, 5 mM CaCl₂, 5 mM Mg(Ac)₂, 0.1 mM EDTA, 10 mM HEPES, 0.1% Triton X-100, 1 mM DTT, 1× protease inhibitor) was added. Cells were collected using a cell scraper and immediately flash-frozen in liquid nitrogen. For chromatin preparation, frozen cells were thawed and crosslinked with 5% paraformaldehyde (PFA) for 10 min at room temperature. Crosslinking was quenched with 125 mM glycine for 5 min, followed by centrifugation at 1,000 × g for 3 min at 4 °C. The supernatant was discarded, and the nuclei pellet was resuspended in 500 µl Nelson buffer (140 mM NaCl, 20 mM EDTA, 50 mM Tris-HCl pH 8.0, 0.5% NP-40, 1% Triton X-100, 1× protease inhibitor). The nuclei suspension was homogenized using an Ultra-Turrax T10 homogenizer (IKA) at power setting 4 for 10 s. The suspension was centrifuged at 10,000 × g for 2 min at 4 °C, and the pellet was washed once with Nelson buffer. The nuclei pellet was then weighed and resuspended in RIPA buffer (140 mM NaCl, 1 mM EDTA, 1% Triton X-100, 0.1% sodium deoxycholate, 10 mM Tris-HCl, 1% SDS, 1× protease inhibitor). Samples were incubated on a rotating wheel at 4 °C for 10 min and then sonicated for 30 cycles (30 s on / 30 s off). To assess chromatin shearing, an aliquot was decrosslinked with RNase A and proteinase K treatment for 1 h at 65 °C. DNA was purified using the Zymo ChIP DNA Clean & Concentrator Kit following the manufacturer’s instructions. Fragment size was analyzed with a Bioanalyzer 2100 (Agilent) using a High Sensitivity DNA Kit, and DNA concentration was measured with a Qubit 2.0 fluorometer (Thermo Fisher Scientific). For immunoprecipitation, 1 µg of chromatin was incubated with 3 µg of antibody against H3K27me3 (Millipore) or an IgG isotype control (Millipore).

## Statistical analysis

Statistical analysis was done using GraphPad Prism version 9. All graphs are represented as mean ± SEM unless stated otherwise. A two-tailed unpaired t-test or one-way ANOVA was used for analysis.

## Data availability

Supplemental tables can be found here: https://nextcloud.dzne.de/index.php/s/LeKaTZNRZzNTpQ3

Supplemental Figures can be found here: https://nextcloud.dzne.de/index.php/s/Ymr23zA2k5NZFSF

## Acknowledgments

AF was supported by the DFG (Deutsche Forschungsgemeinschaft) SFB1286 and GRK2824; The German Federal Ministry of research, Technology and Space (BMFTR) via the ERA-NET Neuron project EPINEURODEVO; The EU Joint Programme-Neurodegenerative Diseases (JPND) – EPI-3E; Germany’s Excellence Strategy - EXC 2067/1 390729940 and FS was supported by the GoBIO project miRassay (16LW0055) by the BMFTR. ID is supported by NIH RF1AG078299. DKV received funding as a fellow of the Hertie Network of Excellence in Clinical Neuroscience.

## Author Contributions

R.P. planned and conducted the majority of experiments and wrote this manuscript. M.S.S. performed nuclei sorting. D.M.K. and T.P. helped with bioinformatic analysis. S.B., A.-L.S., and F.S. orchestrated RNA sequencing. D.K.V. contributed iPSC-derived microglia and ID contributed human brain tissues. A.F. conceptualized the project, supervised progress, and wrote this manuscript. R.P. is part of the IMPRS graduate school for Neuroscience. All authors have read and agreed to the published version of this manuscript.

